# Silicone Oil-Induced Glaucomatous Neurodegeneration in Rhesus Macaques

**DOI:** 10.1101/2022.06.06.494980

**Authors:** Ala Moshiri, Fang Fang, Pei Zhuang, Haoliang Huang, Xue Feng, Liang Li, Roopa Dalal, Yang Hu

## Abstract

Previously, we developed a simple procedure of intracameral injection of silicone oil (SO) into mouse eyes and established the mouse SOHU (SO-induced ocular hypertension under-detected) glaucoma model with reversible intraocular pressure (IOP) elevation and significant glaucomatous neurodegeneration. Because the anatomy of the non-human primate (NHP) visual system closely resembles that of humans, it is the most likely to predict human responses to diseases and therapies. Here we replicated the SOHU glaucoma model in rhesus macaque monkeys. All six animals that we tested showed significant retinal ganglion cell (RGC) death, optic nerve (ON) degeneration, and visual functional deficits at both 3 and 6 months. In contrast to the mouse SOHU model, IOP changed dynamically in these animals, probably due to individual differences in ciliary body tolerance capability. This acute NHP glaucoma model closely recapitulates the major features of glaucomatous neurodegeneration in humans, and is therefore suitable for studying the pathology of primate RGC/ON, assessing experimental therapies for neuroprotection and regeneration, and therefore for translating relevant findings into novel and effective treatments for patients with glaucoma and other neurodegenerations.

## Introduction

Glaucoma, the most common cause of irreversible blindness, is characterized by progressive peripheral to central loss of retinal ganglion cells (RGCs) and their axons in optic nerve (ON) ^1–4^. Although glaucoma can occur at any intraocular pressure (IOP) level ^5^, elevated IOP is associated with accelerated progression ^1–4,6,7^. Lowering IOP is the only available treatment but fails to completely prevent the progression of glaucomatous neurodegeneration ^8–11^. Neuroprotectants that promote RGC/ON survival, transplantation of stem cell-derived RGCs to replace lost RGCs, and regeneration therapies to stimulate RGC soma and axon regrowth are promising neural repair strategies to restore vision in glaucoma patients ^12,13^. To translate exciting laboratory findings into effective neuroprotective and regenerative treatments, pre-clinical testing in a disease-relevant, translation-enabling animal glaucoma model that closely resembles human patients is critically important.

We recently developed a silicone oil (SO)-induced ocular hypertension under-detected (SOHU) glaucoma mouse model ^14–16^ based on the well-documented, SO-induced human secondary glaucoma that occurs as a complication of vitreoretinal surgery ^17,18^. By blocking aqueous flow to the anterior chamber with a single intracameral injection of SO that induces pupillary block, this SO injection causes accumulation of aqueous and significant IOP elevation in the posterior chamber, and subsequent progressive RGC and ON degeneration. Importantly, the ocular hypertension of the SOHU model can be reversed quickly and definitively by easily removing SO from the anterior chamber ^14–16^. However, there is a recognized gap in the translation of successful neuroprotective and regenerative therapies identified in rodent models of glaucoma to treatment for glaucoma patients. Rodents have known limitations that may impede translation of potential therapeutics: differences in immune system responses, ON head (ONH) architecture, and brain structures and circuitry may contribute to differences in pathogenesis between rodents and primates and, therefore, to critically different responses to therapeutics. Despite the many benefits of the mouse SOHU model, a higher experimental animal species is needed for pre-clinical translation research.

The anatomy of the non-human primate (NHP) visual system closely resembles that of humans and includes a similar distribution of rods and cones, a specialized macula and fovea and lamina cribrosa not present in rodent, comparable contrast sensitivity and visual acuity, and almost identical retinocortical architecture ^19,20^. An NHP glaucoma model is the most likely to predict human responses to ocular hypertension and therapies, and the rhesus macaque monkey has been used successfully in experimental glaucoma research ^21,22^. Since SO-induced pupillary block causes secondary glaucoma in both human patients and mice, we reasoned that the same procedure may be adapted to different animal species with minimal confounding factors. Here we report the development of a novel NHP glaucoma model in rhesus macaque monkeys, in which intracameral SO injection causes severe RGC and ON degeneration and visual function deficits. We expect this model to be useful for studying primate RGC pathophysiology, assessing experimental neuroprotective and regenerative therapies, and therefore for translating relevant findings into novel and effective treatments for patients with glaucoma and other neurodegenerations.

## Results

### Intracameral injection of SO in rhesus macaque monkey causes RNFL thinning and decreases PhNR

We injected roughly 100 μl SO into the anterior chamber of the right eyes of 6 macaque monkeys (**Table 1**), filling 80% SO of the anterior chamber with complete covering of the pupil (**Fig. 1A**). Retinal morphology and function were assayed before SO injection and at different time points after. These assays included fundus imaging, spectral-domain optical coherence tomography (SD-OCT), and electroretinography (ERG) (**Fig. 1B**). Thinning of the retina nerve fiber layer (RNFL) measured by OCT is used clinically as a biomarker for RGC/ON degeneration ^23–25^. We measured the RNFL thickness of the animals and detected edema (thickening) of RNFL in the SOHU eyes at 3-month post injection (3mpi) and significant thinning at 6mpi (**Fig. 2A,B**), indicating inner retina neurodegeneration. We also examined the visual function of these macaques. The photopic negative response (PhNR) of the photopic full-field ERG is a negative-going wave that occurs after the b-wave in response to a brief flash and reflects the function of RGCs and their axons in general. Its amplitude is reduced early in human glaucoma ^26^, which also correlates well with structural loss in NHP glaucoma ^27^. Both b-wave and PhNR’s amplitudes decreased in the SOHU eyes at all time points after SO injection, but only reached statistical significance at 1mpi (**Fig. 2C**), suggesting functional deficits of the inner retina.

**Table 1.** Animal Information.

| **ID** | **Sex** | **Date of Birth** | **Weight (Kg)** |
| --- | --- | --- | --- |
| **38361** | **F** | **05/10/2007**<br>**(13yrs)** | **9.68** |
| **42946** | **F** | **05/31/2012**<br>**(8yrs)** | **9.91** |
| **44193** | **M** | **04/06/2014**<br>**(6yrs)** | **9.16** |
| **44639** | **M** | **05/28/2014**<br>**(6yrs)** | **12.95** |
| **44876** | **F** | **05/11/2015**<br>**(6yrs)** | **10.38** |
| **45513** | **M** | **06/22/2015**<br>**(6yrs)** | **8.43** |

**Figure 1.**
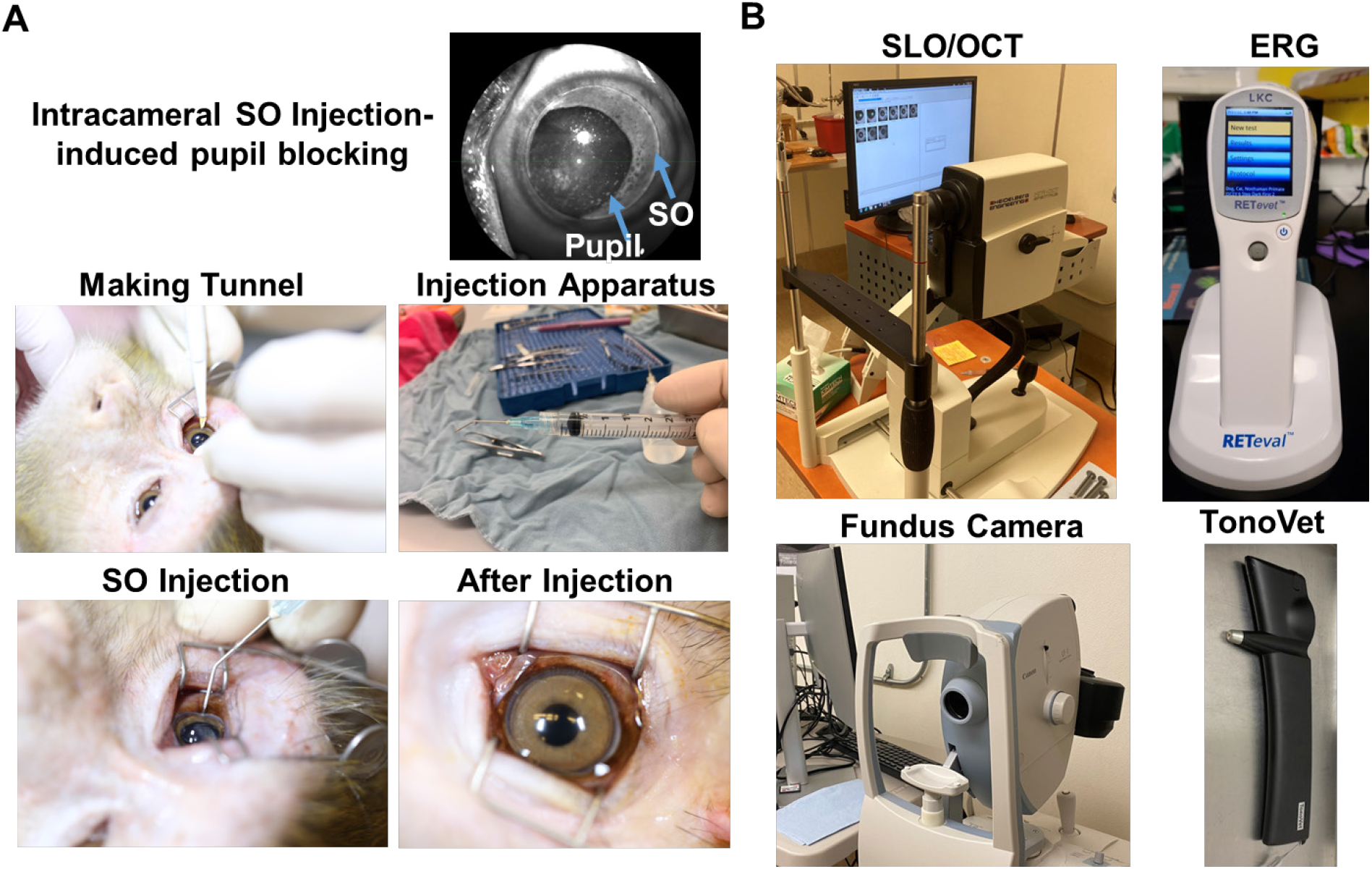
Intracameral SO injection and in vivo assays of rhesus macaque monkey eyes. (**A**) The procedures of SO intracameral injection in monkey eye. (**B**) The equipment used for in vivo assays, including SLO/OCT, ERG, fundus imaging, and TonoVet for IOP measurement.

**Figure 2.**
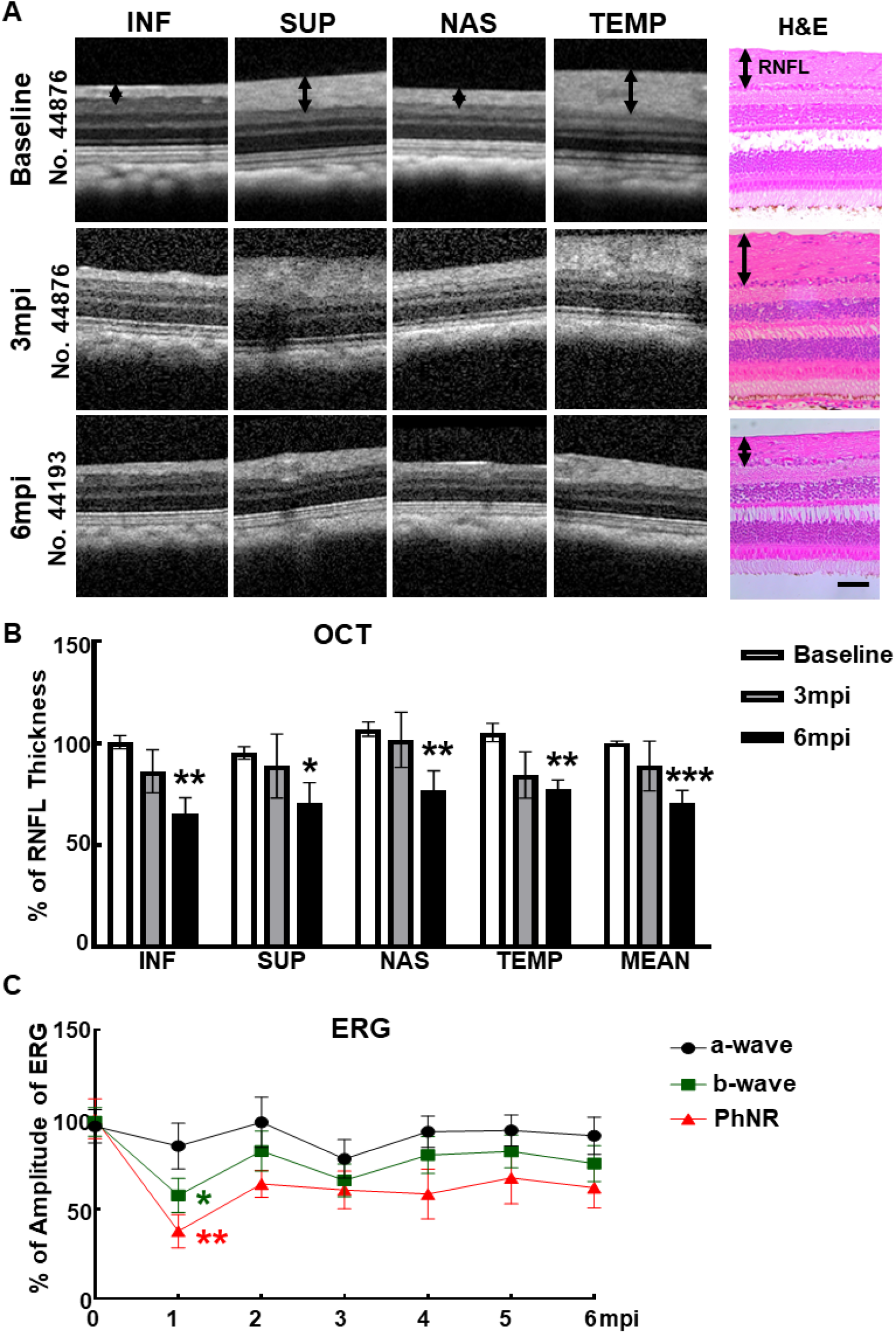
Visual function and morphological deficits of SOHU monkey eyes. (**A**) Longitudinal SD-OCT imaging of SOHU retinas at inferior (I), superior (S), nasal (N), and temple (T) quadrants; and the H&E staining of retina sections. (**B**) Measurements of RNFL thickness at different time points, represented as percentage of SOHU eyes compared to CL eyes. Data are presented as means ± s.e.m, n = 6 for 3mpi and n = 4 for 6mpi, *: *P*<0.05, **: *P*<0.01, ***: *P*<0.001, Student’s t test. (**C**) Longitudinal ERG recording of macaque eyes at different time points after SO injection and the measurements of the amplitudes of a wave, b wave and PhNR, represented as percentage of the amplitudes in the SOHU eyes, compared to the CL eyes. Data are presented as means ± s.e.m, n = 6 for 1-3mpi and n = 4 for 4-6mpi, *: *P*<0.05, **: *P*<0.01, One-way ANOVA with Tukey’s multiple comparison test.

### Significant RGC and ON degeneration of the SOHU eyes at 3mpi and 6mpi in all tested animals

To confirm the glaucomatous neurodegeneration, we euthanized two animals at 3mpi and four animals at 6mpi for histological analysis of post-mortem retina and ON. Consistent with the *in vivo* structural and functional deficits detected in the living animal, retinal wholemounts revealed significant RGC somata loss in the SOHU eye throughout the peripheral to the central retinas at both 3mpi and 6mpi (**Fig. 3A,B**); and semithin cross-sections showed significant RGC axon degeneration in ON at both 3mpi and 6mpi (**Fig. 3C,D**), indicating significant glaucomatous neurodegeneration of the SOHU eyes.

**Figure 3.**
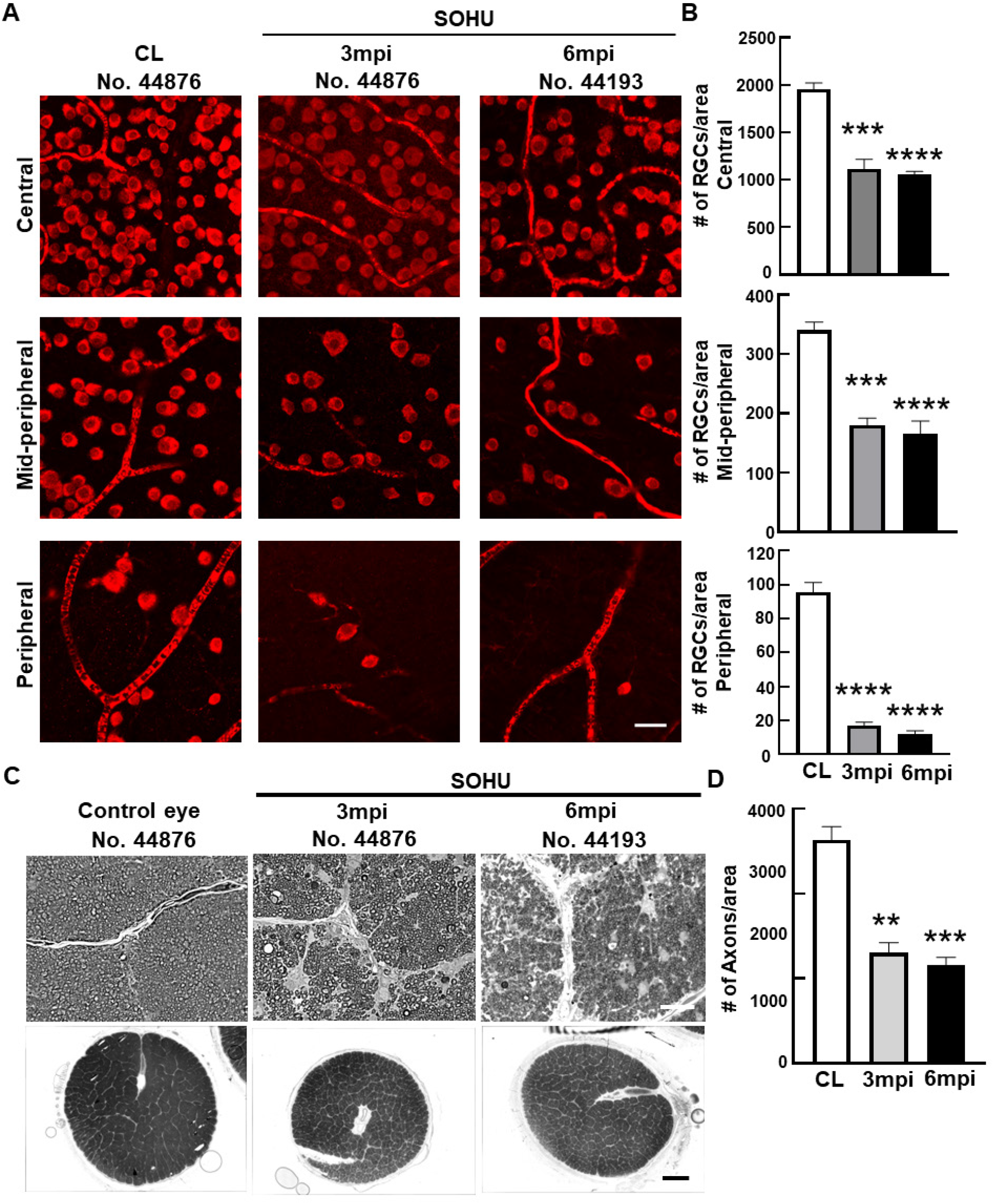
Severe RGC and ON degeneration in SOHU eyes at 3mpi and 6mpi. (**A**) Confocal images of wholemount retinas showing surviving RBPMS-positive (red) RGCs in the peripheral, mid-peripheral, and central retina at 3and 6mpi. Scale bar, 20 μm. (**B**) Quantification of surviving RGCs in the peripheral, mid-peripheral, and central retina. CL: contralateral control eyes. Data are presented as means ± s.e.m, n = 2 for 3mpi and n = 4 for 6mpi, ***: P<0.001, ****: P<0.0001, one-way ANOVA with Tukey’s multiple comparison test. (**C**) Light microscope images of semi-thin transverse sections of ON stained with PPD in the corresponding groups. Upper panel: 100 ×, Scale bar, 20 μm; lower panel: 60 ×, Scale bar, 500 μm. (**D**) Quantification of surviving RGC axons in ON. Data are presented as means ± s.e.m, n = 2 for 3mpi and n = 4 for 6mpi, **: *P*<0.01, ***: P<0.001, One-way ANOVA with Tukey’s multiple comparison test.

### Dynamic IOP changes in the SOHU macaque eyes associated with ciliary body atrophy

Surprisingly, these macaques showed different IOP dynamics after SO injection. In two animals (#44876 and #45513), IOP was elevated immediately after SO injection (15 to 19 mmHg and 13 to 22 mmHg, **Fig. 4A**). Because restrictions of the Primate Center then precluded measuring the IOPs before 1mpi or more frequently than once a month thereafter, we could not measure the IOP earlier or more often. Therefore, we do not know for the duration of the transient IOP elevation after SO injection. However, all six animals showed substantial ocular hypotension at 1mpi and 2mpi: IOPs of the SOHU eyes were much lower than their baselines or their contralateral control eyes (**Fig. 4A,B**). The ocular hypotension lasted from 1mpi to 3mpi in two animals (#44876 and #45513) and from 1mpi to 5mpi in one animal (#44639); IOP returned progressively to normal between 2-6mpi in three animals (#42946, #44639, and #44193) that we maintained for 6 months. In one animal (#38361) IOP was much higher than normal from 3-5mpi, at first fell significantly when SO was removed from the eye at 5mpi, then returned to normal one month later. The sequence of changes in this animal indicated that the SO-induced pupillary blocking was the cause of IOP elevation, and that simply removing the SO reversed the pupillary blocking and ocular hypertension. Because we missed the measurement at the 2mpi time point for this animal (#38361), we assume that the IOP of the SOHU eye recovered from ocular hypotension and became elevated between 1mpi and 3mpi.

**Figure 4.**
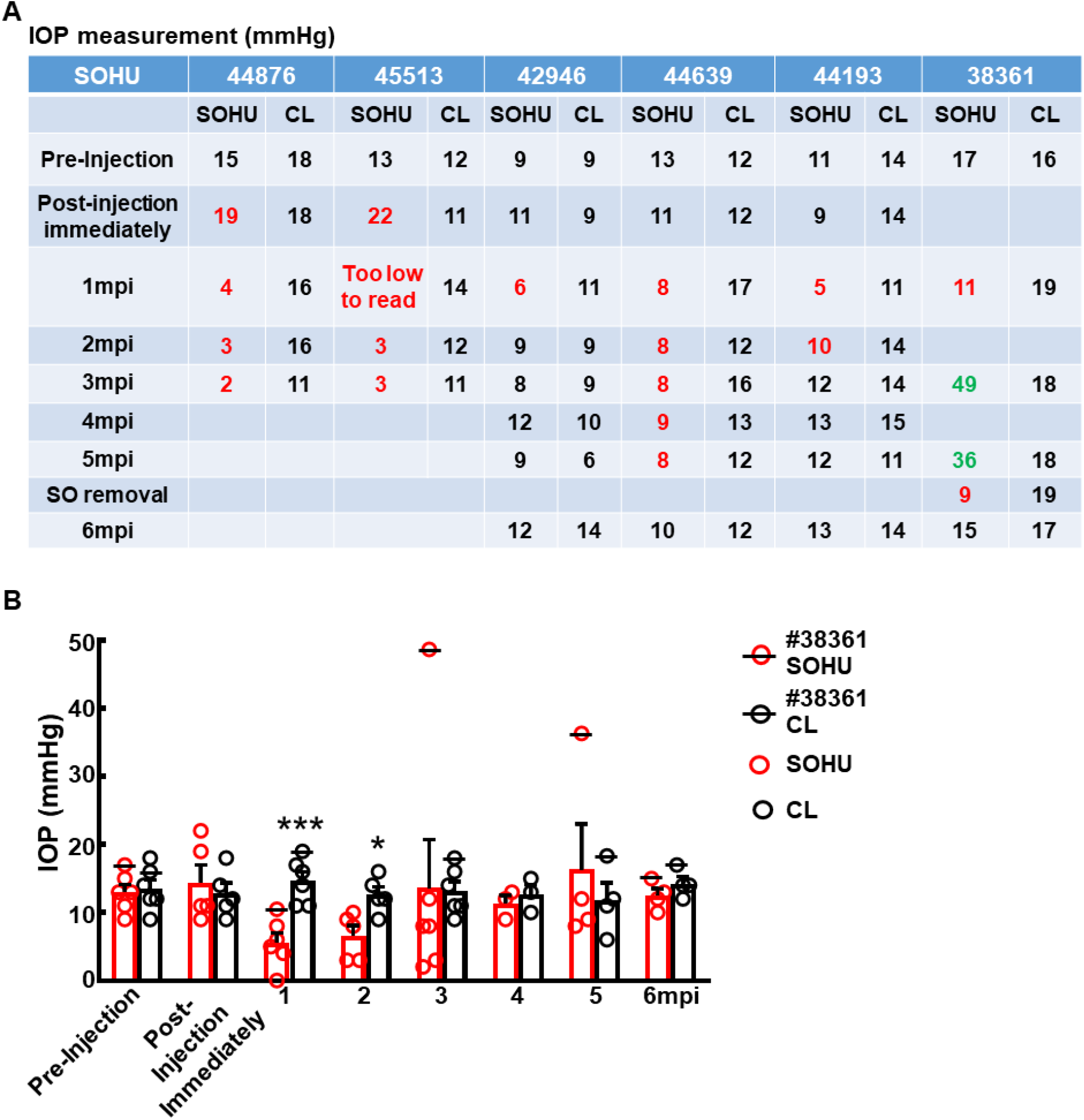
Dynamic IOP changes of SOHU eyes. Table (**A**) and Bar graph (**B**) presentation of longitudinal IOP measurements of experimental (SOHU) eyes and contralateral control (CL) eyes at different time points after SO injection. mpi: month post injection. Data are presented as means ± s.e.m, n = 6 (1-3mpi) and n = 4 (4-6mpi) of each group; *: p<0.05, ***: p<0.001, Student’s t test. Red numbers are ocular hypotension and green numbers are ocular hypertension.

We suspect that the pupillary blockade caused a substantial elevation of the IOP acutely, which led to ciliary body “shutdown”, as in some human patients ^28^. The subsequent lasting ocular hypotony then happened due to ceased aqueous production from ciliary body. Indeed, the ciliary body was severely atrophied in the SOHU eyes of all animals, revealed by H&E staining of the anterior segments of the eyes (**Fig. 5** and **Supplementary Figure 1A**). There was no inflammation or obvious deformation of cornea, sclera, iris, or lens, although the pupils of the SOHU eyes were fixed in the mid-dilated state (**Supplementary Figure 1B**), suggesting a transient high IOP elevation, which may result in ischemic iris sphincter muscle and consequently limitation in constriction, as in patients with acute angle closure glaucoma ^29^.

**Figure 5.**
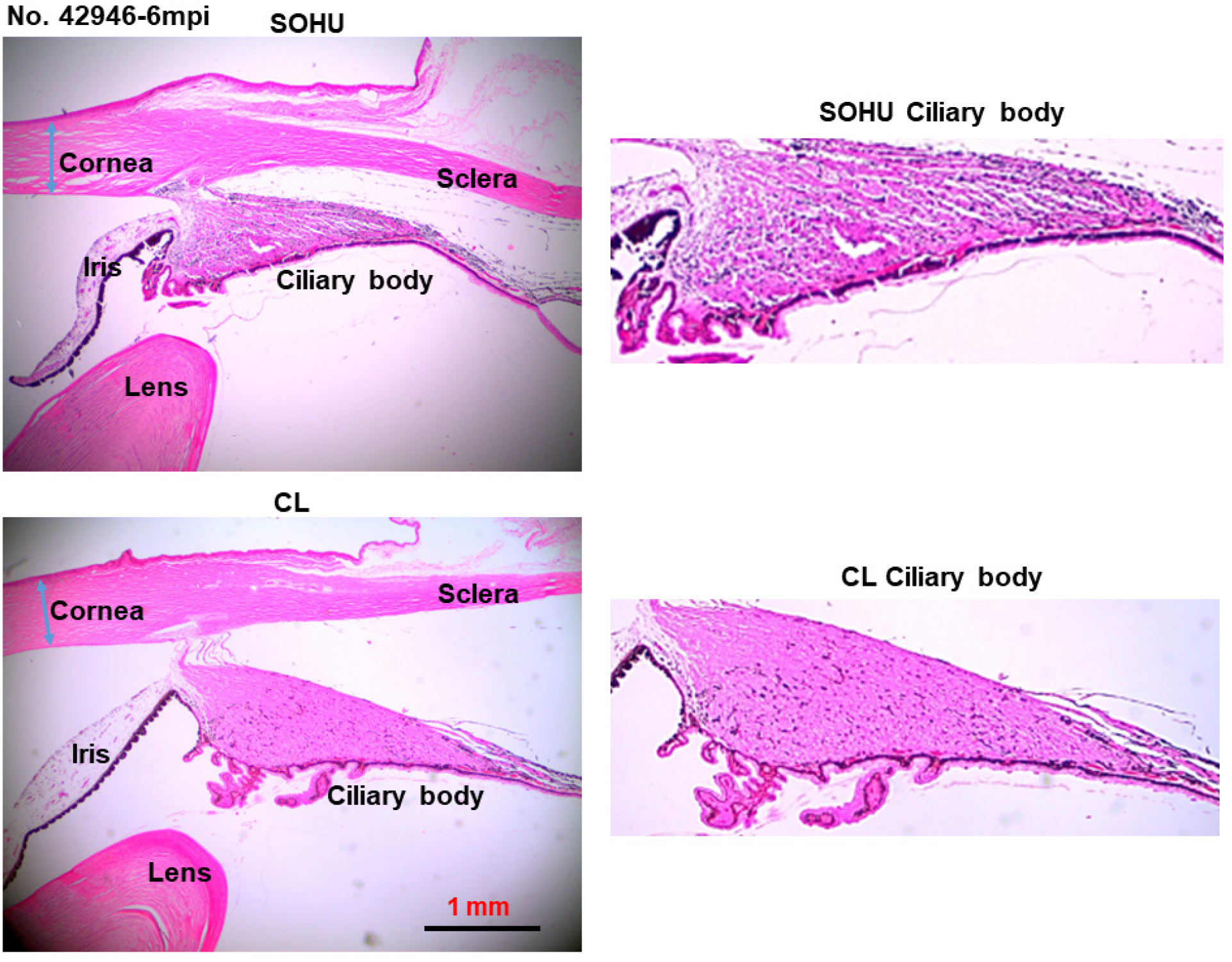
Ciliary body atrophy in SOHU eyes at 6mpi. Anterior chamber sections stained with H&E and imaged with 2× lens; and enlarged images of ciliary body, showing loose arrangement, larger interfibrous areas, and increased cellular invasion in muscle fibers.

### ON head “cupping” is present in the SOHU eye with persistent IOP elevation

A characteristic morphological feature of human glaucoma is enlargement of the depression in the center of the ONH, called glaucomatous “cupping”^30,31^. Strikingly, live fundus imaging with confocal scanning laser ophthalmoscopy (cSLO) readily detected this signature morphological change of glaucoma in the SOHU macaque eye (#38361) by at 3 and 5mpi (**Fig. 6A**), corresponding to IOP elevation (**Fig. 4**). That ONH cupping is absent in the mouse SOHU model further confirms the similarity between macaque and human eyes. This characteristic glaucomatous optic cup enlargement was even more obvious in OCT live imaging by radial B-scan centered through the ONH (**Fig. 6B**). Based on previously developed measurement of the anatomic features of the macaque ONH ^31,32^, we applied the Visualization Toolkit (VTK) to reconstruct and delineate the OCT imaging data (**Supplementary Figure 2A**). We used inner limiting membrane (ILM), Bruch’s membrane opening (BMO), the two discrete points at either side of the neural canal, and the BMO reference plane as references to acquire minimum rim width (MRW), rim volume (RimV), and cup volume (CupV). Obvious shortening of MRW, shrinking of RimV, and enlarging of CupV were detected in the SOHU eye compared to contralateral control eye (**Supplementary Figure 2B**). The H&E staining of the ONH confirmed the “cupping” phenotype and significant thinning of RNFL (**Supplementary Figure 2C-E**). The lamina cribrosa is a trabecular connective tissue to support RGC axons at the ONH ^33^. Its deformation, such as increased curve and depth, may correlate with RNFL thinning in glaucoma patients ^34,35^. Interestingly, collagen staining of the ONH of the SOHU eye also showed lamina cribrosa bowing (**Supplementary Figure 2D**). ONH “cupping” cannot be found by fundus SLO images or OCT images in the eyes of the other macaques without persistent IOP elevation (**Supplementary Figure 3A,B**), indicating the correlation of prolonged ocular hypertension and ONH “cupping”.

**Figure 6.**
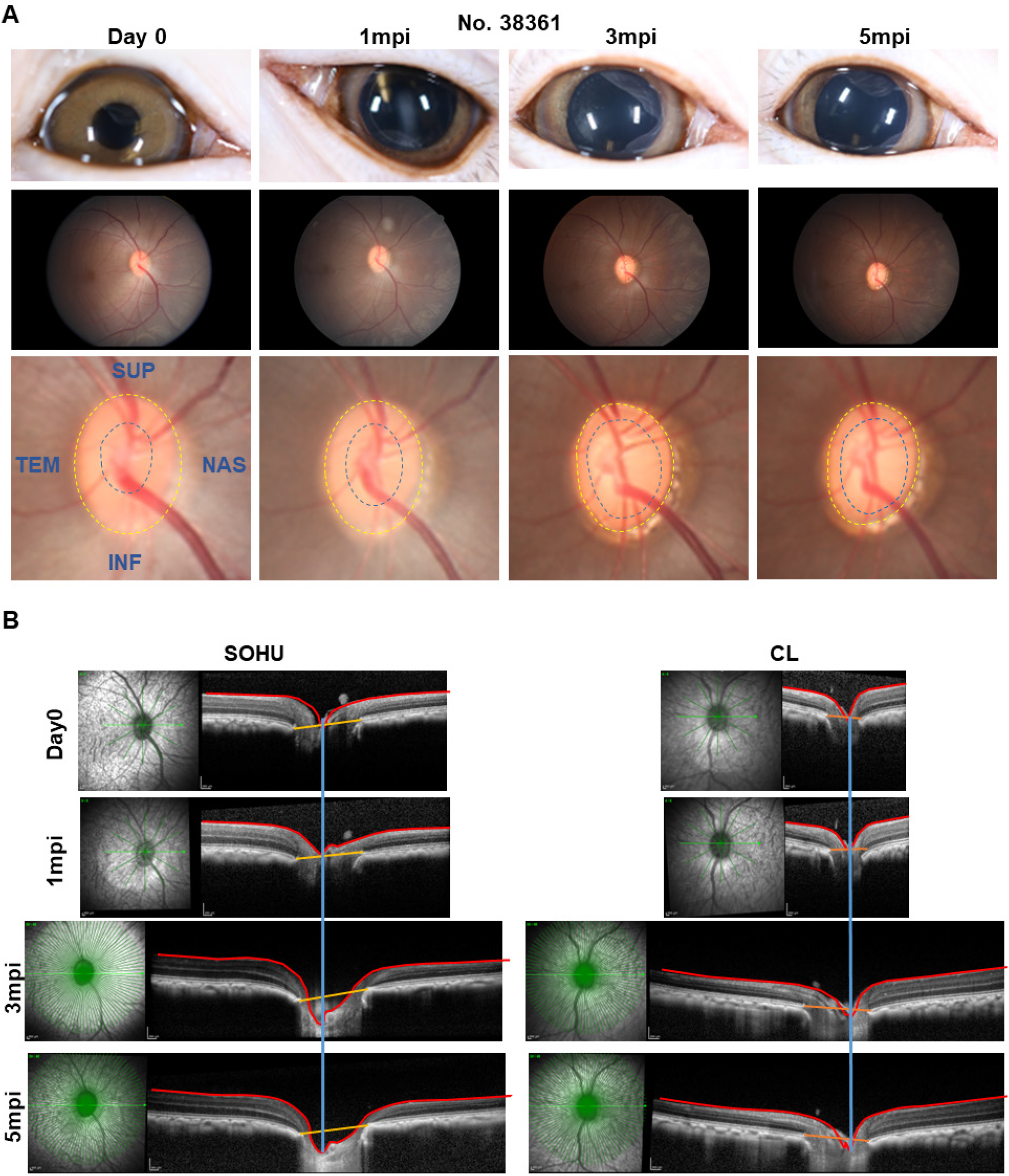
ONH “cupping” in animal #38361 associated with IOP elevation. (**A**) The retinal fundus images of the SOHU eye before and after SO injection. Yellow dotted line outlines the optic disc; blue dotted line outlines optic cup. (**B**) Longitudinal SD-OCT imaging of macaque ON head with 48 radial B-scans acquired over a 30° area at 768 A-scans per B-scan, ART=16 repetitions.

## Discussion

The present report establishes a straightforward and minimally invasive procedure, a single intracameral injection of SO, to induce reproducible glaucomatous RGC and ON degeneration within 3-6 months in rhesus macaque monkeys. The model mimics acute secondary glaucoma caused by pupillary blocking and can be used to study the pathogenesis of neurodegeneration and to select urgently needed neuroprotectants and regeneration therapies that are unrelated to IOP management. Within 3-6 months of a simple SO intracameral injection, the SOHU eyes of all monkeys studied showed a highly consistent array of findings: significant thinning of RNFL, decreased visual function (PhNR), and loss of RGC somata and axons. The reversible intracameral SO injection does not cause overt anterior ocular structural damage other than the ciliary body while simulating acute glaucomatous RGC and ON changes. Therefore, this inducible, reproducible, and clinically relevant NHP neurodegeneration model can be used to decipher the molecular mechanisms of transient ocular hypertension-induced glaucomatous degeneration in primate, and to preclinically assess the efficacy and safety of experimental strategies for neuroprotection and regeneration.

A unique feature of this NHP model is the transient IOP elevation-induced ciliary body “shock”. Unlike mouse, but as can happen in humans ^28^, the NHP ciliary body seems very vulnerable to acutely elevated IOP, which first caused it to stop generating aqueous humor and ocular hypotension, and ultimately leads to atrophy. All six monkeys that we tested consistently developed persistent intraocular hypotension and histological evidence of ciliary body atrophy, although we captured the initial transient IOP elevation before ocular hypotension in only two animals. Unfortunately, most animals (five out of six monkeys) studied did not fully recover normal ciliary body function. Ciliary body function appeared to recover in part, however, since they became able to maintain low or normal IOP in the presence of SO-induced pupillary blocking within the time period of the experiment (3-6 months). Despite the absence of long-lasting chronic ocular hypertension, all five animals showed similar RGC and ON degeneration as the one animal with persistent ocular hypertension. This suggests that transient acute IOP elevation causes the neurodegeneration. From our mouse study ^16^, we learned that although SO removal allows IOP to return quickly to normal, it does not stop the progression of glaucomatous neurodegeneration in the SOHU model. This result is also consistent with the clinical observation that visual field loss can progress aggressively in some glaucoma patients whose IOP is maintained at a relatively low level. Thus, this NHP SOHU model can be used to determine the efficacy of experimental neuroprotection treatment when IOP is low after an initial period of pathogenic ocular hypertension, simulating clinical IOP treatment. Advanced retinal imaging and visual function assays that are available for humans can be applied to this primate glaucoma model. These assays will identify morphological and functional changes in RGCs and ON that can serve as potential biomarkers in glaucoma and other optic neuropathies. Since optic neuropathy can also be associated with other central nervous system (CNS) neurodegenerative diseases ^36^, including multiple sclerosis ^23,37,38^, Alzheimer’s disease ^39,40^, and amyotrophic lateral sclerosis ^41,42^, this model may be broadly applicable to diverse CNS degenerative diseases.

One animal (#38361) was able to recover rather quickly from ciliary body shock and resume adequate aqueous humor production, which increased IOP due to pupillary blocking. It is notable that the characteristic glaucomatous ONH “cupping” was associated with persistent ocular hypertension in this animal but absent from the other animals without persistent IOP elevation.

Ocular vascular dysfunction has long been known to be correlated with the incidence of glaucoma and acutely elevated IOP in patients with angle-closure glaucoma, and secondary glaucoma can induce central retinal artery occlusion with ischemic damage of the inner retina ^43–46^. We previously detected ocular ischemia with inner retina damage and outer retina sparing in the severe variant of the SOHU mouse model ^16^, consistent with findings in rats with acutely elevated IOP ^47–50^. Interestingly, we also detected branch retinal artery occlusion in the animal (#38361) with elevated IOP, demonstrating another similarity between the ocular hypertension in NHP and the human acute glaucoma-related syndrome. We do not know what causes the variable ciliary body responses of different animals. Age may play a role since #38361 was much older (13yrs) than the other five animals (6-8yrs); the middle-aged ciliary body in this animal may be more resilient than younger ciliary bodies. Further systematic studies with additional senior, middle-aged, and young NHP animals are needed to clarify the reasons and to further optimize this model. For example, a modified SOHU model like the one that we developed in mouse that induces and maintains a moderate elevation of IOP through frequent pupil dilation ^16^ may prevent the acute severe IOP elevation causing ciliary body shock.

## Methods

### Animal

The animals in this study were rhesus macaques (Macaca mulatta) born and maintained at the California National Primate Research Center (CNPRC). The CNPRC is accredited by the Association for Assessment and Accreditation of Laboratory Animal Care (AAALAC) International. Guidelines of the Association for Research in Vision and Ophthalmology Statement for the Use of Animals in Ophthalmic and Vision Research were followed. All aspects of this study were in accordance with the National Institutes of Health (NIH) Guide for the Care and Use of Laboratory Animals and all methods are reported in accordance with ARRIVE guidelines. Phenotyping and ophthalmic examinations were performed according to an animal protocol approved by the University of California Davis Institutional Animal Care and Use Committee and Stanford University School of Medicine.

### Intracameral injection of SO

The procedure is similar to the published protocol ^14,15^ but with modification for monkey eyes. Sedation was achieved by intramuscular injection of ketamine hydrochloride (5-30 mg/kg IM) and dexmedetomidine (0.05-0.075 mg/kg IM). The eyes were prepped in a usual sterile fashion for ophthalmic surgery including topical anesthetic 0.5% proparacaine hydrochloride (Akorn, Somerset, New Jersey) followed by 5% betadine to the ocular surface and adnexa. A disposable 15-degree blade was used to make a side-port incision at the corneal limbus to enter the anterior chamber inferiorly near the 6 o’clock position in order to minimize the likelihood of oil leaking out of the eye. Silicone oil (SO, 1,000 mPa.s, Silikon, Alcon Laboratories, Fort Worth, Texas) in a 3 cc syringe on a bent 25 gauge cannula was introduced into the anterior chamber. SO was injected little by little, stopping intermittently with gentle pressure applied to the posterior aspect of the limbal incision to allow for aqueous humor to exit the eye.

Oil was injected to fill the anterior chamber to a physiologic depth with roughly ~70-80%silicone oil and to cover the entire pupil with ~ 100 μl volume. After the injection, the wound was tested to insure it was self-sealing and veterinary antibiotic ointment (BNP Ophthalmic Ointment, Vetropolycin, Dechra, Overland Park, Kansas) was applied to the surface of the injected eye. The contralateral control eyes received a mock injection with no penetration of the eye. Animals were monitored by a trained technician and a veterinarian at all times.

### Removing SO from the anterior chamber

The procedure is similar to the published protocol ^14,15^ with modification for monkey eyes. Briefly, after the animal was anesthetized the eye was prepped in a sterile fashion as above. A superior (12 o’clock) corneal side-port incision was made using a 15-degree blade at the corneal limbus. A 3 cc syringe filled with sterile balanced salt solution (BSS Plus, Alcon Laboratories, Ft. Worth, Texas) with a 25 gauge bent cannula was introduced into the anterior chamber and saline was gently injected little by little while periodically allowing oil to egress from the same incision by gently applying pressure to the posterior aspect of the wound. After removing all of the oil and replacing it incrementally with BSS to a physiologic depth, the cannula was removed and the wound was checked to be self-sealing, after which antibiotic ointment was applied.

### Eye examinations and retinal fundus imaging

Sedated ophthalmic examination included measurement of intraocular pressure (IOP) using rebound tonometry (Icare TA01i, Finland) while the animal was held upright and with careful attention not to apply any pressure to the globe. Three IOP measurements were taken and averaged at each exam date. Examination also included pupillary light reflex testing, external and portable slit lamp examination, as well as dilated (Tropicamide 1%, Phenylephrine 2.5%, Cyclopentolate 1%) indirect ophthalmoscopy. Sedation was achieved by intramuscular injection of ketamine hydrochloride (5-30 mg/kg IM) and dexmedetomidine (0.05-0.075 mg/kg IM). Animals were monitored by a trained technician and a veterinarian at all times. Color and red-free fundus photographs were obtained with the CF-1 Retinal Camera with a 50° wide angle lens (Canon, Tokyo, Japan).

### Spectral-domain optical coherence tomography (SD-OCT) imaging

SD-OCT with confocal scanning laser ophthalmoscopy (cSLO) was also performed (Spectralis® HRA+OCT, Heidelberg, Germany). High-resolution radial and circumferential scans centered on the optic nerve were obtained using a corneal curvature (K) value of 6.5 mm radius. For the high-resolution radial scans of the optic nerve head (ONH), 48 radial B-scans were acquired by 870 nm SD-OCT (Spectralis; Heidelberg Engineering, GmbH), over a 30° area, and 768 A-scans per B-scan at ART=16 repetitions. All repetitive scans were acquired using eye-tracking and averaged to reduce speckle noise. We read in all the images and measured MRW, RimV, and CupV using R program. The codes that we used to calculate MRW, RimV and CupV are at Github (https://github.com/HuLab-Code/ONHV). For each monkey eye, the center of the ONH was estimated and registered during the first imaging session and used to align all follow-up images. All imaging was done by the same ophthalmic imaging team. All OCT images were taken through the center of the pupil. Speculums were used and corneal hydration was maintained through application of topical lubrication (Genteal artificial tears) approximately every 1-2 minutes during imaging sessions. The en-face retinal images were captured with the Heidelberg Spectralis SLO/OCT system equipped with an 870nm infrared wavelength light source and a 30° lens (Heidelberg Engineering). The average thickness of retinal nerve fiber layer (RNFL) around the optic nerve head was measured manually with the aid of Heidelberg software. The investigators who measured the thickness of RNFL were masked to the treatment of the samples.

### Electroretinography (ERG) recording

After dilation, a full-field ERG (ffERG) containing six different tests was performed on each eye following a 30-minute dark adaptation period. ERG-Jet electrodes (item #95-011) were coupled with the RETeval instrument (LKC Technologies, Gaithersburg, MD, United States), as previously described ^51^. A standard flash electroretinogram was performed according to the approved protocol of the International Society for Clinical Electrophysiology of Vision (ISCEV). There were four dark adapted tests (0.01 cd*s/m2, 3.0 cd*s/m2, 10.0 cd*s/m2, and oscillatory potentials 3.0 cd*s/m2). After 10 minutes of light adaptation, two additional tests were performed (3.0 cd*s/m2 single flash with measurement of the photopic negative response and photopic flicker 3.0 cd*s/m2). Both time (ms) and amplitude (μV) were obtained for each test on each eye. Single flash tests measured an a-wave and b-wave. Oscillatory potentials measured five wave points and a sum. In the photopic flicker test, the first wave point is reported. Measurements were recorded and displayed using the manufacturer’s software.

### Immunohistochemistry of whole-mount retina and RGC counting

The detailed procedure has been published before ^14,15,52^ with modification to accommodate large monkey eyes. Briefly, after intravitreal injection with 10% formalin in PBS, the eyes and optic nerves were dissected out, post-fixed with 10% formalin for 24 hours at room temperature. Retinas were dissected out and washed extensively in PBS before blocking in staining buffer (10% normal goat serum, Sigma-Aldrich, and 2% Triton X-100 in PBS) for half an hour. RBPMS guinea pig antibody made at ProSci Inc (Poway, California) according to publications ^53,54^ was diluted (1:4000) in the same staining buffer. Floating retinas were incubated with primary antibodies overnight at 4°C and washed 3 times for 30 minutes each with PBS. Secondary antibodies (Cy3) were then applied (1:200; Jackson ImmunoResearch, West Grove, Pennsylvania) and incubated for 1 hour at room temperature. Retinas were again washed 3 times for 30 minutes each with PBS before a cover slip was attached with Fluoromount-G (SouthernBiotech, Birmingham, Alabama). For RGC counting, whole-mount retinas were immunostained with the RBPMS antibody, 6 fields sampled from each region (periphery, mid-periphery, and center retinas) using a 20× lens with Keyence epifluorescence microscope, and RBPMS^+^ RGCs of each image (540 μm × 720 μm) were counted manually with Fiji/ImageJ. The investigators who counted the cells were masked to the treatment of the samples.

### ON semi-thin sections and quantification of surviving axons

The detailed procedure has been published before ^14,15,52^. Briefly, the ON was exposed by removing the brain and post-fixed *in situ* using 2% glutaraldehyde/ 2% PFA in 0.1M PB for 4 hours on ice. Samples were then washed with 0.1M PB 3 times, 10 minutes each wash. The ONs were then carefully dissected out and rinsed with 0.1M PB 3 times, 10 minutes each wash. They were then incubated in 1% osmium tetroxide in 0.1M PB for 1 hour at room temperature followed by washing with 0.1M PB for 10 minutes and water for 5 minutes. ONs were next dehydrated through graded ethanol, infiltrated in propylene oxide and epoxy, and embedded in epoxy at 60°C for 24 hours. Semi-thin sections (1 μm) were cut on an ultramicrotome (EM UC7, Leica) and collected 2 mm distal to the eye. The semi-thin sections were attached to glass slides and stained with 1% para-phenylenediamine (PPD) in methanol: isopropanol (1:1) for 35 minutes. After rinsing 3 times with methanol: isopropanol (1:1), coverslips were applied with Permount Mounting Medium (Electron Microscopy Sciences, Hatfield, Pennsylvania). PPD stains all myelin sheaths, but darkly stains the axoplasm only of degenerating axons, which allows us to differentiate surviving axons from degenerating axons ^55^. The whole ON were imaged with a 100x lens of a Keyence fluorescence microscopy to cover the entire area of the ON without overlap. Four areas of 108 μm × 144 μm were cropped, and the surviving axons within the designated areas counted manually with Fiji/ImageJ. After counting all the images taken from a single nerve, the mean of the surviving axon number was calculated for each ON. The investigators who counted the axons were masked to the treatment of the samples.

### Anterior segments and retina cross sections and H&E and Trichrome Staining

Monkey eyes were enucleated and immediately fixed in 10% formalin for 36 hours at room temperature. They were processed through graded alcohol and xylene, then infiltrated and embedded in paraffin. Six-micron sections were taken and stained with Hematoxylin & Eosin (H&E) to look at the cell nuclei, extracellular matrix, and cytoplasm using Nikon Eclipse (E800) microscope. Standard protocol was followed to stain these slides. The Trichrome kit was purchased from Abcam (ab 150686) to study collagenous connective tissue in sections. Slides were deparaffinized and incubated in preheated Bouin’s fluid for an hour and rinsed in water. They were then incubated in Weigert’s Iron Hematoxylin for 5 minutes, rinsed in water again and then incubated in Biebrich Scarlet/Acid Fuchsin solution for 15 minutes. They were rinsed in water again. Sections were then differentiated in phosphotungstic acid solution for 10-15 minutes (or until collagen is not red), incubated in Aniline Blue solution for 5-10 minutes and rinsed in water. Acetic acid solution was applied to these sections for 3-5 minutes, and slides were then dehydrated in alcohol, cleared in xylene, and mounted with CytoSeal 60 (from Electron Microscopy Sciences, 18006). This stain shows a stronger collagen stain (blue green stain) in glaucomatous eye than the control eye.

### Statistical analyses

GraphPad Prism 7 was used to generate graphs and for statistical analyses. Data are presented as means ± s.e.m. Student’s t-test was used for two groups comparison and One-way ANOVA with post hoc test was used for multiple comparisons.

## Data availability

All data generated or analyzed during this study are included in this published article (and its Supplementary Information files).

## Acknowledgements

We thank Drs. Jeffrey Goldberg, Jie Zhang, Xin Duan, Derek Welsbie, Anna La Torre Vila, and Alan Tessler for critical discussion. Y.H. is supported by CNPRC Pilot Grant, NIH grants EY031063, EY024932, EY023295, EY028106, and EY032518, and grants from Glaucoma Research Foundation (CFC3), BrightFocus Foundation, Chan Zuckerberg Initiative Neurodegeneration Collaborative Pairs Pilot Projects, Stanford SPARK program, and Stanford Center for Optic Disc Drusen. We are grateful for an unrestricted grant from Research to Prevent Blindness and NEI P30 EY026877 to the Department of Ophthalmology, Stanford University.

## Author contributions

Y.H., A.M., and F.F. designed the experiments. A.M. performed surgeries and eye exams, F.F., P.Z., H.H., X.F., L.L., and R.D. processed the samples. Y.H., F.F., and A.M. analyzed the data and prepared the manuscript.

