## Supplementary Figures for "Silicone Oil-Induced Glaucomatous Neurodegeneration in Rhesus Macaques"

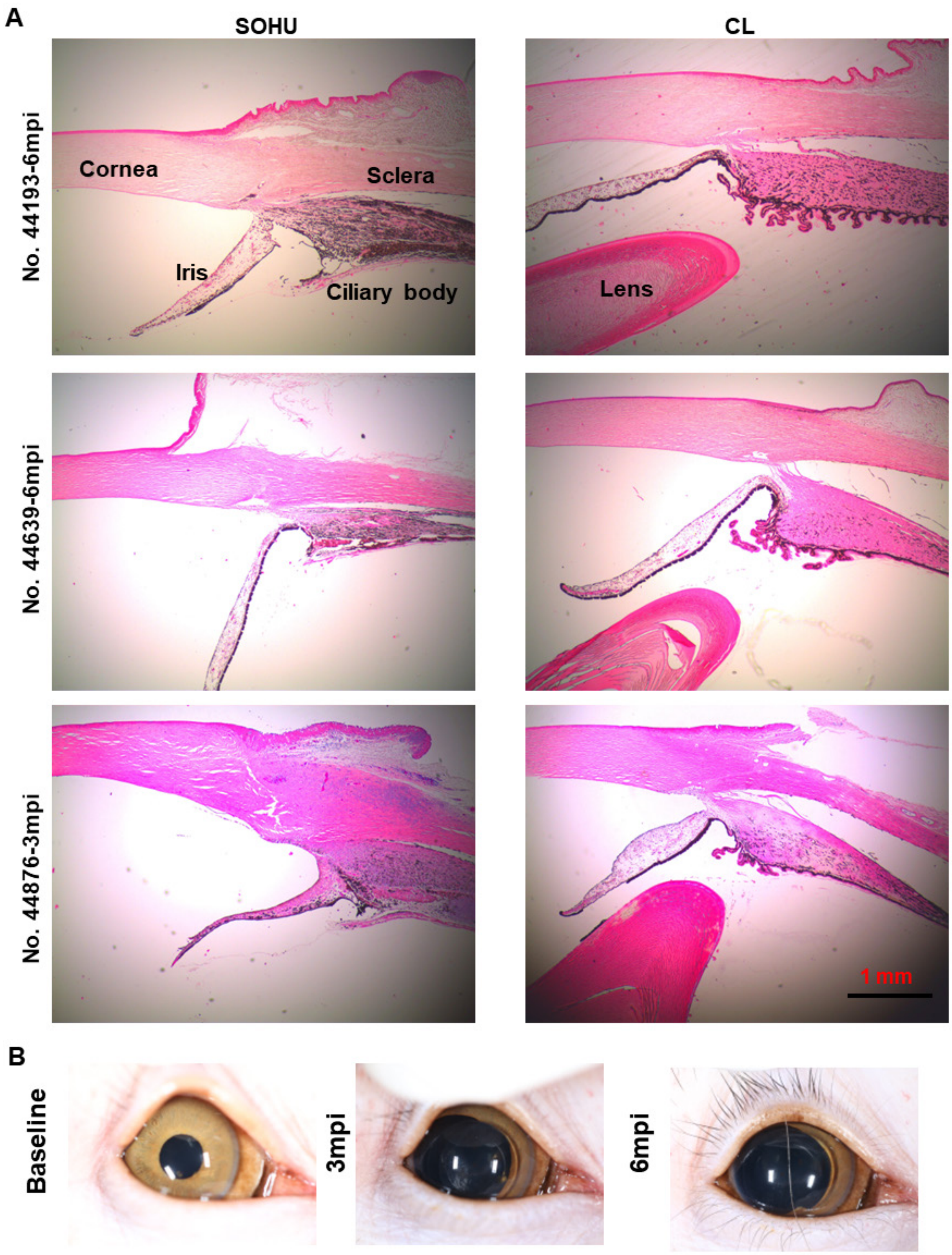


**Supplementary Figure 1. Ciliary body atrophy and dilated pupil in SOHU eyes at 6mpi and 3mpi.** (**A**) Anterior chamber sections stained with H&E and imaged with 2x lens. (**B**) Pupil dilation after SO injection.

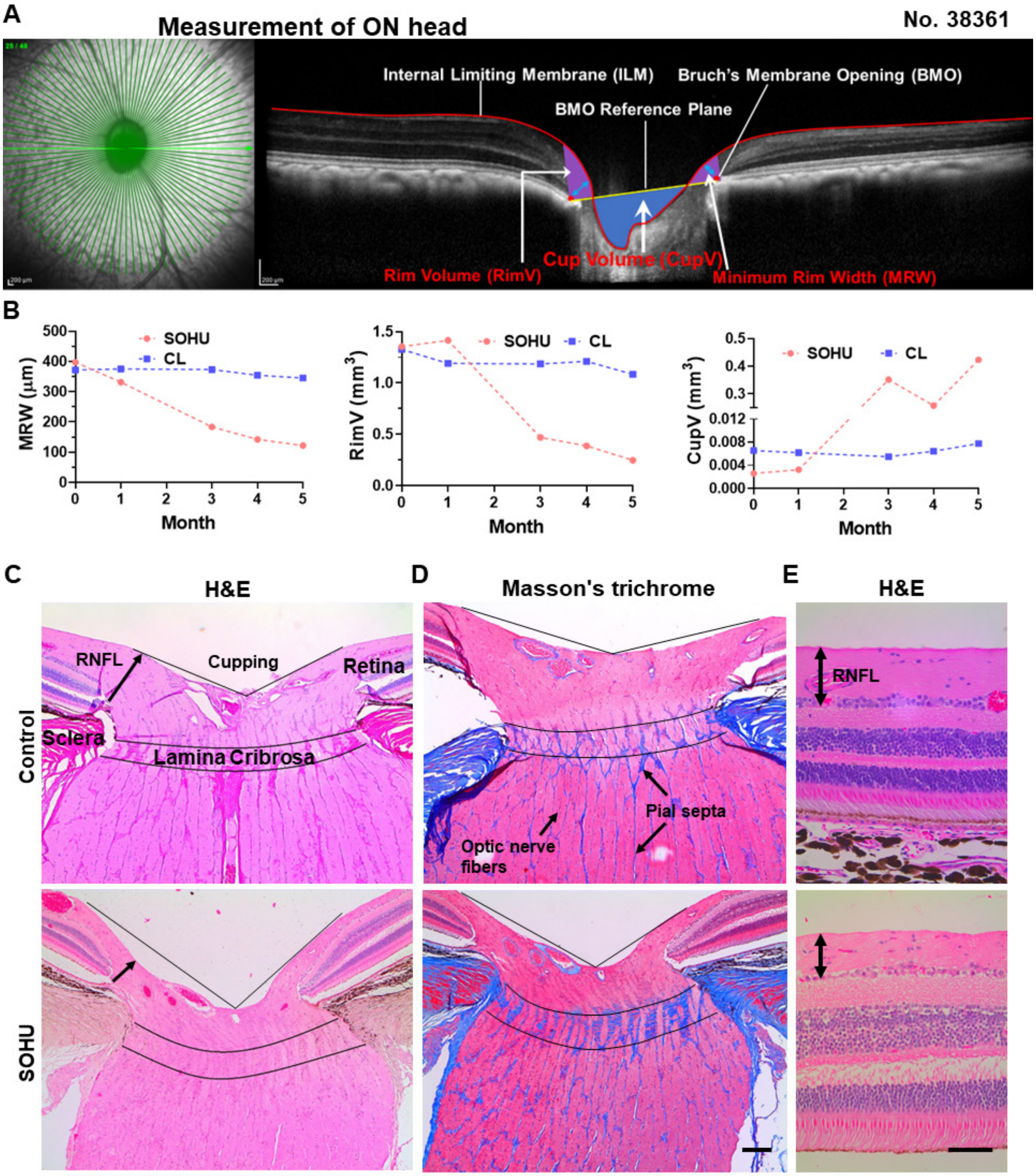


**Supplementary Figure 2. Measurements of ONH "cupping" of animal #38361.** (**A**) Neural tissue parameters: Minimum rim width (MRW, blue double headed arrow) was measured at each delineated Bruch's membrane opening point (BMO, red dot) as the minimum distance to ILM; Rim volume (RimV, purple area) was calculated from the volume bounded by ILM (red line), BMO reference plane (yellow line) and perpendicular line through the BMO; Cup volume (CupV, blue area) was generated from the volume between ILM surface and the BMO reference plane. (**B**) The raw OCT images were transformed to the gray value matrix. The noise from the image acquisition and transformation was reduced by Gaussian filter. After the denoise processing, then 3D model and the MRW, RimV and CupV were established by Otsu's thresholding method and convolutional neural network method. (**C**) ONH sections stained with H&E. (**D**) ONH sections stained with Masson's trichrome. Scale bar = 200 μm. (**E**) Retina sections stained with H&E. Scale bar = 50 μm.

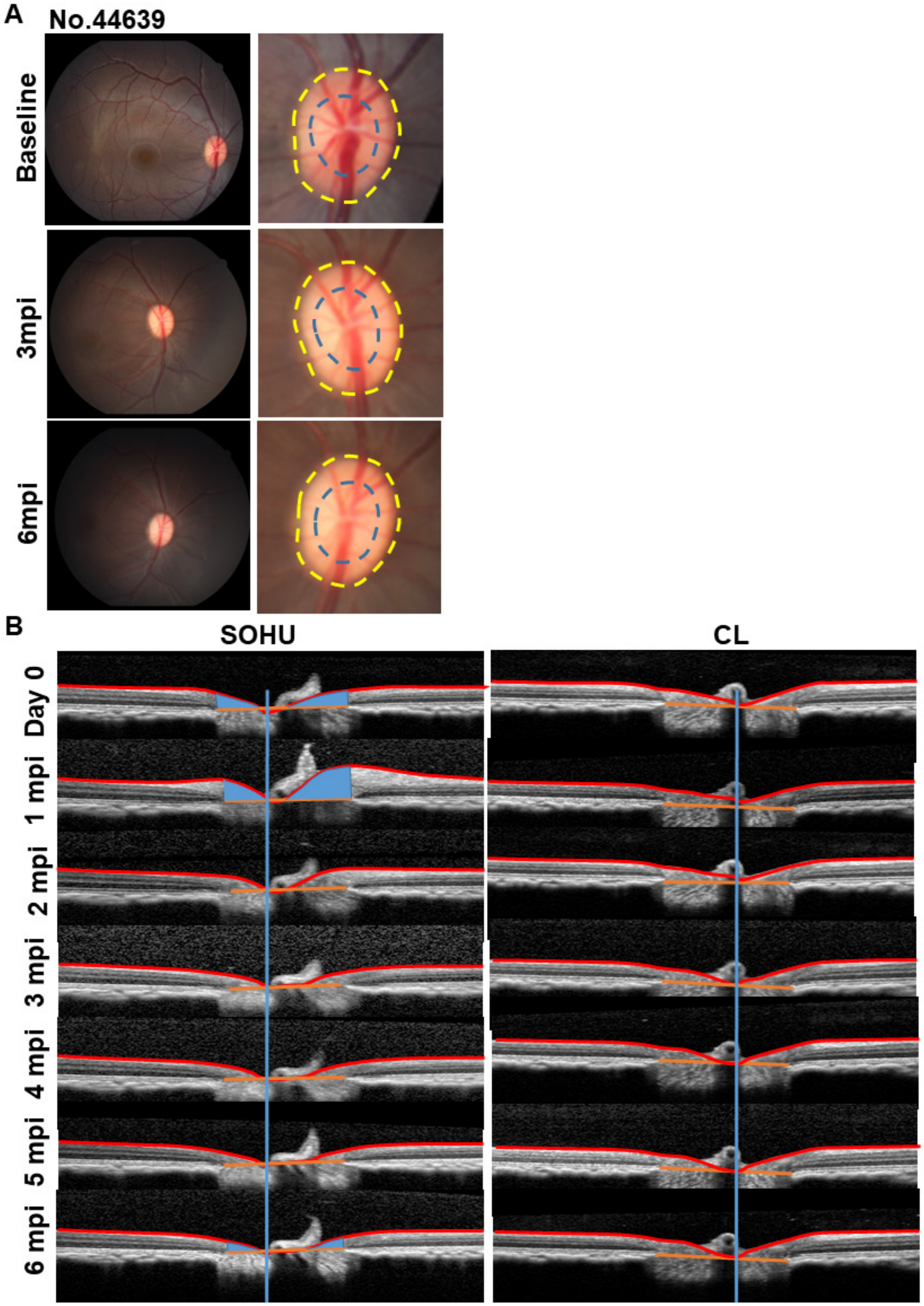


**Supplementary Figure 3. No detectable ONH "cupping" in the other five animals.** (**A**) The retinal fundus images of the SOHU eye before and after SO injection. Yellow dotted line outlines the optic disc; blue dotted line outlines optic cup. (**B**) Longitudinal SD-OCT imaging of ON heads.
